# Widespread Negative Frequency-Dependent Selection Maintains Diversity in the Legume-Rhizobia Symbiosis: Balancing nodulation may explain the paradox of rhizobium diversity

**DOI:** 10.1101/153866

**Authors:** Eleanor Siler, Maren L. Friesen

## Abstract

The evolutionary origin and ecological maintenance of biodiversity is a central problem in biology. For diversity to be stable through time, each genotype or species must have an advantage when rare. This negative frequency-dependence prevents deterministic extinction and mitigates the stochastic loss of diversity (*1–4*). However, models of mutualism typically generate positive frequency-dependence that reduces diversity (*5–8*). Here, we report empirical evidence for negative frequency-dependence in the legume-rhizobium mutualism within a single host generation, a phenomenon that we term balancing nodulation. Balancing nodulation increases rare rhizobia across all 13 legume genera investigated to date, at high and low inoculum densities, and with minimal genetic differentiation between rhizobia strains. While the mechanism generating this phenomenon is currently unknown, balancing nodulation could actively maintain variation in the rhizobia-legume symbiosis.

---

Biological diversity is rapidly declining (*9*), yet biodiversity underlies ecosystem functioning and resilience *(*10–12*).* Mutualists provide key resources and services, including pollination, symbiotic nitrogen fixation, and digestion (*13*). These beneficial interactions are predicted to be degraded under global change *(*14*)* and their ability to respond to changing environments requires genetic variation (*15*). Thus, understanding the evolutionary and ecological maintenance of diversity is a central and urgent problem in biology (*11,16*), particularly mutualisms. Theory delineates the conditions for diversity maintenance under antagonistic interactions (*1,2,17)* and antagonistic coevolution is predicted to drive rapid coevolution through the Red Queen mechanism (*18*). Antagonistic coevolution can create and maintain diversity via negative frequency-dependent selection (*8,19*). Antagonistic coevolution can readily generate negative frequency-dependence across generations, such as reciprocal shifts in *Linum marginale* resistance alleles and *Melampsora lini* virulence alleles (*20*) and the rapid increase of rare MHC alleles in experimental stickleback populations (*21*). In contrast, models of mutualistic interactions typically generate positive frequency dependent selection, which acts against the maintenance of diversity (*8,19, 22*) and has been observed in plant mutualisms with arbuscular mycorrhizal fungi (*23*). The model symbiosis between legumes and rhizobia, in which soil bacteria colonize host roots and fix atmospheric nitrogen in root nodules, is crucial to agriculture and nitrogen cycling (*24, 25*). Contrary to theoretical expectations, rhizobia contain large amounts of genetic (*26*) and functional (*27*) diversity. While symbiont diversity has been proposed to arise through antagonistic coevolution driven by cheaters, evidence for this is weak in rhizobia and other systems (*28,29*). This supports a model of coordinated coevolution in mutualisms, further intensifying the paradox of rhizobial diversity.

For symbiont diversity to be actively maintained, strains must have a fitness advantage when rare that they lose when they become common (*2*). Negative frequency-dependent selection means that each strain can invade a community when rare, preventing deterministic extinction and opposing diversity loss due to stochastic sampling. Frequency dependence is quantified by measuring the fitness of two competing strains across a variety of starting ratios. In the simplest scenario, with two strains A and B that are equally competitive and without frequency dependence, the strains ratio in the inoculum (*I_A_/I_B_*) equals the ratio of the strains nodule occupancy (*N_A_/N_B_*). A log-log plot of the nodule versus inoculum ratios has a slope of 1. If one strain is more competitive by a factor *C*, then the log-ratio of the strains in nodules will be

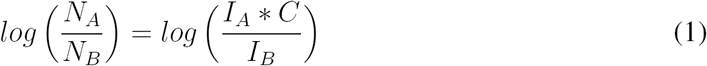

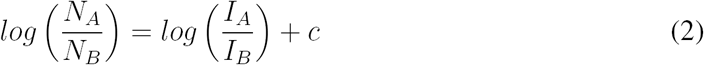

where *c = log(C).* Thus, when two strains of differing competitiveness are inoculated at varying ratios and the resulting nodule ratio is plotted on a log-log scale, the data are linear with slope 1 and intercept c. Under this scenario, the more competitive strain will always out-compete the other strain, driving it to extinction (Fig. 1A). Frequency-dependent selection is introduced by adding the frequency-dependence coefficient *k*, such that

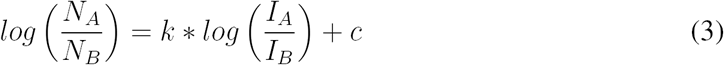

Equation 2 was introduced to measure relative rhizobia competitiveness (*c*) and is widely used in agronomic studies *(*30*).* When *k* differs from 1, strain competitiveness is frequency dependent. This has dramatic consequences for diversity, which can be illustrated by the difference equation that describes the strain ratio dynamics between iterations of plant growth:

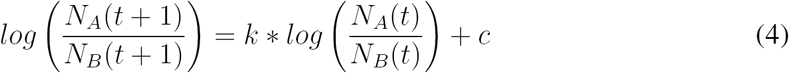

which assumes for simplicity that the inoculum ratio in the next generation is the strain nodule ratio in the previous generation. Typically, a single rhizobium cell initiates each root nodule and multiplies to 10^5^ – 10^8^ cells (*31*), making nodule occupancy an adequate proxy for rhizobia fitness. Equation 3 results in the equilibrium strain ratio *Log(N_A_/N_B_*) = c/(1 – *k*); this equilibrium is dynamically stable when *k* < 1 and unstable when *k* > 1 (Appendix 1). When *k* > 1 the more common strain has an advantage, thwarting diversity and resulting in a monomorphic population (Fig. 1B). When *k* < 1, the rarer strain has an advantage, actively maintaining symbiont diversity (Fig. 1C). We note that the presence of multiple cells founding nodules, known as mixed nodules, does not impact strain frequency dependence (Appendix 2).

**Figure 1:**
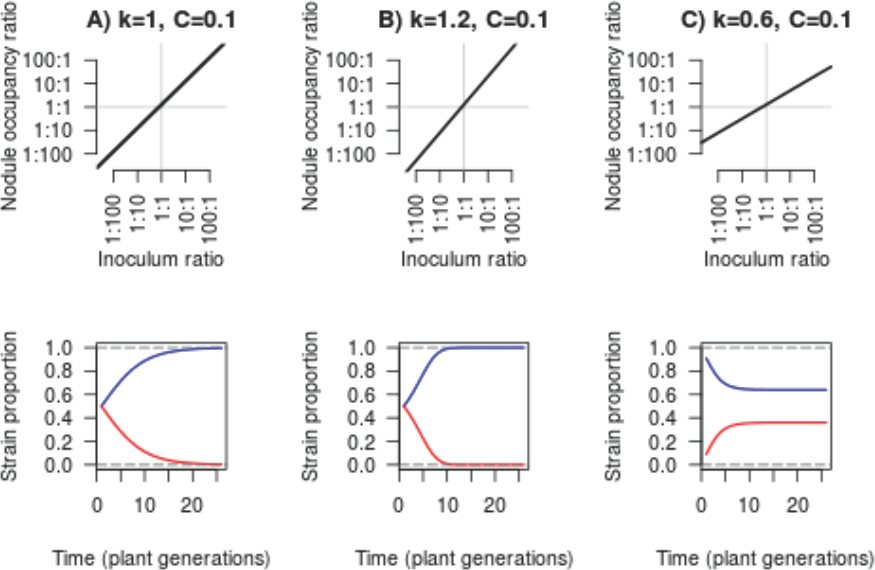
Results from theoretical rhizobial competitions (row 1) and their effects on longterm population dynamics (row 2). Blue: strain A; red: strain B. Column 1: no frequency dependence, column 2: positive frequency dependence, and column 3: negative frequency dependence.

To determine the prevalence of frequency-dependent selection in the rhizobia-legume symbiosis, we performed a meta-analysis of 110 experiments from 30 publications in which rhizobial competition was measured at multiple inoculum ratios (Supplemental Data 1; Supplemental Methods). Using equation 2, we determined the frequency-dependence coefficient (*k*) for each experiment. We found substantial negative frequency-dependent selection: the frequency-dependence coefficient *k* has a mean of 0.56 (95 percent confidence interval: 0.46 – 0.66), far less than the null expectation of *k* =1 (Fig. 2A). This effect is large enough to have considerable ecological implications. For two equally competitive rhizobia strains, a strain that comprises 10 percent of the inoculum will form approximately 30 percent of the nodules, and when either strain comprises 1 percent of the inoculum it will form about 9 percent of the nodules (Fig. 2B). Systematic experimental error, such as poor inoculum preparation, is perhaps the simplest explanation for this phenomenon. However, given that 30 publications across several labs – and decades – contain evidence of balancing nodulation, systematic experimental error seems incapable of consistently producing such large differences between stated inoculum density and actual nodule occupancy (Fig. 2B). We found no evidence of publication bias, perhaps because the experiments were designed to find the competition coefficient, *C*, not the *k* value (Fig. S1). Hence, our data unequivocally demonstrate that rhizobia typically experience an advantage when rare during nodulation of a host legume, a phenomenon that we term balancing nodulation after balancing selection, which maintains alleles at a stable polymorphism (*21*). Balancing nodulation occurs within a single host generation and thus could be caused by ecological processes favoring rare strains during rhizosphere colonization or strain-dependent regulation of nodulation by the host plant.

**Figure 2:**
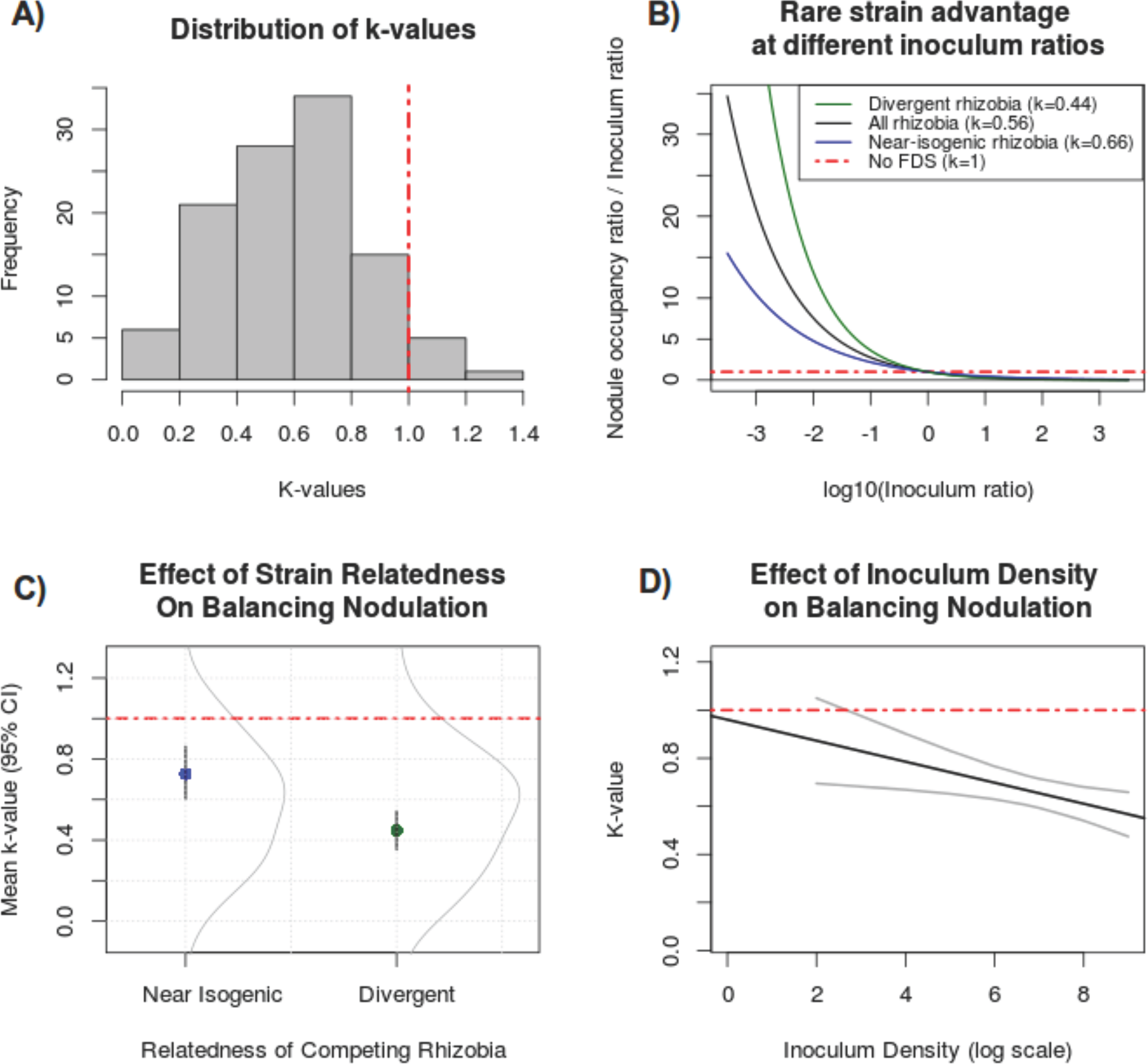
A) Balancing nodulation in the legume-rhizobia symbiosis. The distribution of k-values (gray) compared to the null hypothesis of no frequency dependence (red line). B) Relative over-representation of rhizobia at different inoculum frequencies based on our metaanalysis results. C)Effect of strain relatedness on balancing nodulation. Dots: mean value, black lines: 95 percent confidence intervals, gray curves: distributions of k-values. D) Effect of inoculum density on balancing nodulation. Gray lines: 95 percent confidence interval.

We identified two covariates that affect the strength of balancing nodulation: strain relatedness (near-isogenic vs. divergent strains) and inoculum density (Table 1, Fig. 2C, Fig. 2D). Our finding that divergent rhizobia experience stronger balancing nodulation supports the hypothesis that this phenomenon arises from ecological divergence or strain-specific plant responses. Surprisingly, even near-isogenic rhizobia strains, differing only by a genetic marker, demonstrate balancing nodulation (*k* = 0.66). We also found that as the inoculum density increases, balancing nodulation becomes stronger (Table 1, Fig. 2D). This is consistent with the hypothesis that rhizobia competition contributes to balancing nodulation, although there is no relationship between the strength of balancing nodulation and the difference in strain competitiveness (Fig. S3).

**Table 1:** The effects of covariates on frequency dependent selection, using publication as a random effect.

| **Model Covariate** | **Mean Effect** | **95% CI** | **AIC** | **BIC** | **Significant?** | **Effect** |
| --- | --- | --- | --- | --- | --- | --- |
| Near-Isogenic | 0.279 | [0.15, 0.40] | -38.2 | -27.4 | Yes | Yes |
| None | 0.56 | [0.46, 0.66] | -27.8 | -17.7 | Yes | Yes |
| Log10(Inoculum Density) | -0.06 | [-0.10, -0.01] | NA | NA | Yes | Yes |
| Sterile | 0.1 | [-0.05, -0.26] | -24.3 | -13.5 | No | Inconclusive |
| Determinate | 0.11 | [-0.08, 0.30] | -24.3 | -13.5 | No | Inconclusive |
| Points Measured | -0.01 | [-0.04, 0.02] | -20 | -9.26 | No | Inconclusive |
| Year | 0.01 | [0.0, 0.01] | -18.3 | -7.55 | No | No |
| Competitiveness | 0 | [-0.019, 0.019] | -18.3 | -7.5 | No | No |
| Marker Type | Non-significant | NA | -11.7 | 9.87 | No | Inconclusive |
| Plant Genus | Non-significant | NA | -1.06 | 39.5 | No | Inconclusive |

Balancing nodulation mediated by the plant would require legumes to acquire, integrate, and respond to information identifying their potential rhizobial symbionts. Although plants do use Nod factors and effectors to differentiate between rhizobial strains, there is no evidence that these pathways operate in a frequency-dependent manner nor are they known to have the specificity to discriminate between near-isogenic strain pairs. In animal systems, several evolutionarily independent adaptive immune systems have been described that all generate negative frequency dependence within a host (*32*). In vertebrates, there are two origins of diverse lymphocyte receptors generated by recombination of either immunoglobulin or leucine-rich repeat gene segments (*32*). Insects lack this recombinational mechanism by use alternative slicing to generate diverse immunoglobulin receptors (*32*). In plants, the highly diverse LRR gene family has been argued as evidence for the absence of an adaptive immune system (*32*), yet there is growing evidence for the potential importance of multiple mobile signals of infection that could convey information regarding microbial identity (*33*). We speculate that balancing nodulation could occur through bacterial surface molecules or diffusible signals that are sensed by the plant and acted upon in a systemic manner.

We emphasize that balancing nodulation does not produce selection that would maintain rhizobium cooperation. In fact, this phenomenon provides one potential explanation for the prevalence of ineffective strains in nature. Many experiments in our dataset demonstrate that balancing nodulation favors ineffective rhizobia over effective ones when the ineffective strains became rare (Supplemental Data 1). Thus, balancing nodulation is distinct from host sanctions (*34*) and partner choice (*35*), both of which promote cooperative symbiont behavior.

Balancing nodulation runs rampant throughout legumes, including Papilionoid and Mimosoid taxa (Fig. 3). All 13 legume genera and all legume species investigated thus far show evidence of balancing nodulation (Fig. S2). The prevalence of balancing nodulation suggests that it confers a broad evolutionary advantage or results from an evolutionarily conserved mechanism. If either legumes or rhizobia experience selection to limit the number of nodules formed in a strain-dependent manner, this would favor the evolution of balancing nodulation. Rhizobia populations in the rhizosphere are several orders of magnitude greater than the number of rhizobia that ultimately form nodules (*31*); it may thus be advantageous for strains to limit their infection of the root to avoid overwhelming their host plant. However, an even larger rhizobial benefit could be had by limiting the infection of unrelated competing strains. From the plant perspective, limiting nodulation of specific strains rather than simply limiting the total number of nodules would not necessarily yield an advantage. However, plants could benefit from having diverse symbionts in two ways. First, associating with multiple strains could provide synergistic benefits if the strains are complementary. However, there are counter-examples in which plants inoculated with two strains perform worse than singly-inoculated plants (*36, 37*). Alternatively, plants could favor rare rhizobia as a form of bet-hedging – symbiont diversity could provide insurance against being overtaken by cheaters or otherwise non-beneficial strains in situations when signals do not accurately predict rhizobia partner quality, or when partner quality depends on environmental context. Finally, balancing nodulation could be a pleiotropic effect of some other unknown plant regulatory mechanism that is strongly selected for, such as controlling pathogen infections.

**Figure 3:**
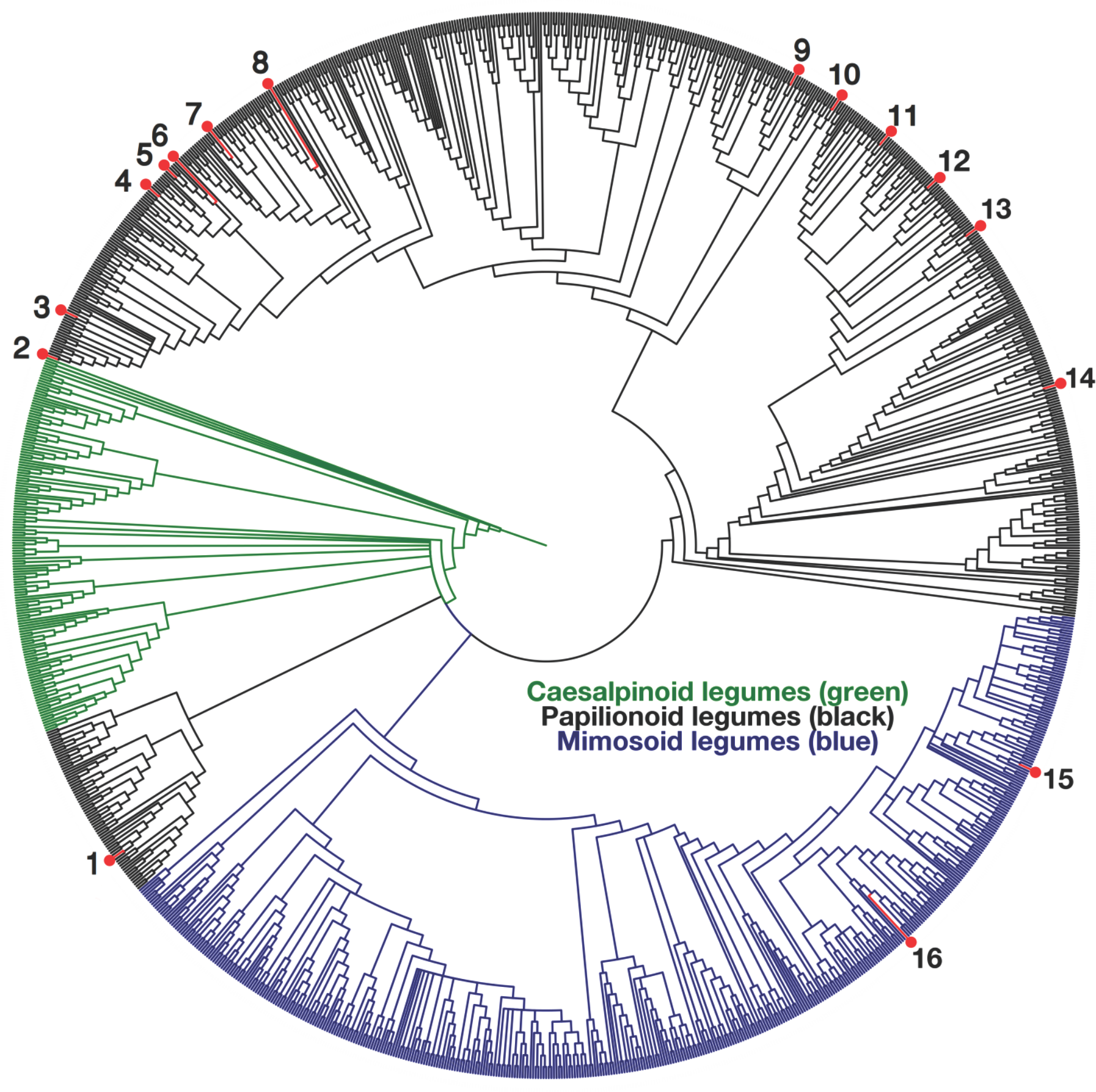
Legume species exhibiting balancing nodulation (red) mapped onto the legume phy-logeny. Species in which balancing nodulation has been detected are 1) *Stylosanthes guia-nenses* 2) *Medicago truncatula* 3) *Medicago sativa* 4) *Trifulium repens* 5) *Trifoliumpratense* 6) *Trifolium subterraneum* 7) *Viciafaba* 8) *Pisum sativum* 9) *Lutus pedunculatus* 10) *Cyamopsis tetragonoloba* 11) *Phaseolus vulgaris* 12) *Macroptillium atropurpureum* 13) *Vigna unguiculata* 14) *Glycine max* 15) *Acacia senegal* and 16) Prosopsis sp.

Balancing nodulation is a ubiquitous phenomenon across legumes, suggesting there is an evolutionary advantage for plants to increase the diversity of their microbial mutualists. While the underlying mechanism is currently unknown, understanding how plants maintain symbiont diversity will be crucial to managing microbial biodiversity for optimal agricultural and planetary health (*38*).

## Acknowledgments

We acknowledge support from NSF DEB 1354878 and NSFIOS 1342793 to MLF. Comments from AW Bowsher, CA Friel, CN Jack, A Garoutte, JS Norman, and S Rowe improved this manuscript.

